# Symbiont spatial organisation is dynamically regulated within cnidarian host tissues

**DOI:** 10.64898/2026.08.28.743919

**Authors:** Alexandra Jilani, Edward S. Allgeyer, Xinyi Li, Minhong Guo, Duygu S. Sevilgen, Alice Ball, Fengzhu Xiong, Susannah B.P. McLaren

## Abstract

The symbiosis with photosynthetic dinoflagellate algae enables corals to build and sustain reef ecosystems. Individual coral polyps hold algal symbionts in their epithelial endoderm cells and lose them under environmental stress, leading to coral bleaching. How the host integrates symbionts into its body plan is not well understood. Here, using a combination of high-resolution imaging, quantitative analysis, and environmental perturbations in the sea anemone *Exaiptasia diaphana* (Aiptasia) and reef-building coral *Pocillopora damicornis*, we uncover a spatial organisation of symbionts along the aboral-oral axis of cnidarian polyps that emerges under the long-range translocation of symbionts between host cells through a fluid-filled cavity. The symbiont distribution becomes specifically enriched in the tentacle bud endoderm during Aiptasia polyp morphogenesis. This pattern can form in darkness and with algae-sized inert spheres, suggesting an innate host-intrinsic mechanism. Symbiont-occupied host cells are mechanically constrained within the endoderm and thus unable to rearrange; instead, they go through cycles of symbiont expulsion and re-uptake via the host gastric cavity, with regionally biased rates of these behaviours providing a route to enrich symbionts in the tentacles. Symbiont organisation is remodelled under increased light in adult coral polyps, with a characteristic pattern of reduced tentacle enrichment, lateral clustering and retention in the body column emerging over a timescale of days. Together, our findings reveal that the spatial organisation of symbionts is dynamically regulated in cnidarian host tissues, a capacity that may shape both the establishment of symbiosis and its resilience under environmental change.

## Introduction

Symbioses between multicellular animals and protist microorganisms enable them to thrive in challenging environments, creating the foundations of new ecosystems. This is exemplified by the partnership between cnidarians, such as corals and sea anemones, and photosynthetic algae. Algal symbionts are internalised into cells in an epithelium that lines the gastric cavity of the host and provide nutrients produced by photosynthesis in exchange for protection and access to host metabolic products^1^. This enables corals to thrive in nutrient-poor seas and build reefs currently supporting a third of all marine species on Earth^2^. However, integrating algal symbionts into an epithelial host tissue context is a non-trivial challenge.

Symbionts are acquired during host development and are carried through a dramatic shape transformation that establishes the characteristic polyp morphology of the host and its orientation within the environment^3,4^. Polyps have a body organised radially-symmetrically around an aboral-oral axis with a two-tissue layered body architecture in which an external ectoderm wraps around a layer of endoderm that lines a fluid-filled gastric cavity^5^. This cavity connects the orally located tentacles that extend towards the light incident from above with the body column that extends aborally and attaches to the underlying substrate (Figure 1A)^5^. Symbionts are phagocytosed from the cavity into the endoderm cells^4^. In the related non-symbiotic sea anemone, *Nematostella vectensis* (Nematostella), polyp morphogenesis is accompanied by extensive remodelling of epithelial tissues^6,7^. In symbiotic cnidarians however, morphogenesis occurs in the presence of relatively large algal symbionts, posing a potential challenge to tissue remodelling with cells occupied by symbionts visibly bulging out of plane of the epithelium^3^. How the host integrates symbionts into its tissue as it acquires its adult morphology is yet to be determined.

**Figure 1.**
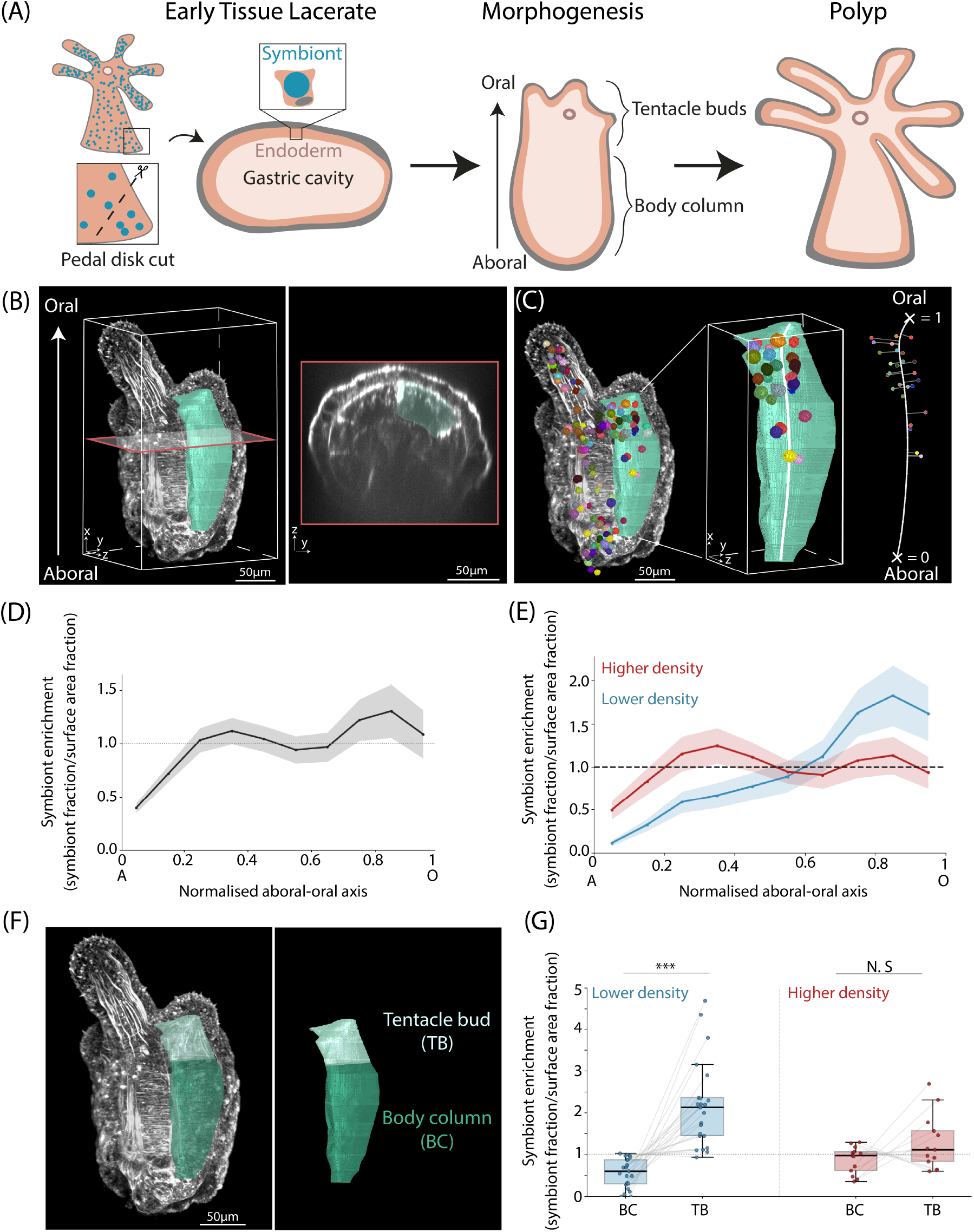
A spatial organisation of symbionts emerges during host morphogenesis. (A) Aiptasia tissue lacerate system to investigate symbiont organisation during host morphogenesis. Tissue lacerates cut from the pedal disk of adult Aiptasia polyps form a fluid-filled gastric cavity and undergo morphogenesis in the presence of symbionts, developing an aboral-oral axis with tentacle buds located at the oral end and ultimately giving rise to a new polyp. (B-C) Analysis workflow for investigating symbiont spatial organisation within the host. (B) Host segments are masked in 3D to obtain a simplified morphological system for investigating symbiont organisation along the aboral-oral axis. A cross-sectional view in the plane indicated on the left panel shows the internal host morphology and actin boundaries used to define the segment. (C) Deep learning-based (Cellpose) segmentation of symbionts is used to obtain symbiont centroids in tentacle budding-stage lacerates. A spline (shown in white) is fit through the centre of each host segment, running from the aboral base to oral tip, and symbiont positions are mapped onto this spline. (D) Pooled symbiont enrichment (binned symbiont proportion/surface area proportion) along the normalised aboral-oral axis (n=36 segments from 18 lacerates, 2114 symbionts). (E) Pooled symbiont enrichment along the normalised aboral-oral axis in low density (blue) and higher density (red) segments (n=23 low segments from 14 lacerates, 522 symbionts; n=13 higher density segments from 9 lacerates, 1592 symbionts). (F) Actin organisation is used to distinguish between the body column and tentacle bud. (G) Symbiont enrichment in the body column (BC) vs tentacle bud (TB) region of segments (blue= low density segments, n=23 segments, p = 7.2×10^-7^; red=higher density segments, n=13 segments, p = 0.27). Lines connect BC and TB pairs for each segment.

Acquired symbionts may be lost post-development. When algal symbionts are exposed to light intensities or temperatures outside their optimal range, their photosystems become stressed, leading to photoinhibition and the production of reactive oxygen species (ROS) that are thought to promote symbiosis breakdown through damage to host cells and decreased nutrient transfer that triggers symbiont expulsion and coral bleaching^8–10^. Interestingly, not all symbioses break down under stress - some persist under unknown mechanisms of adaptation^11–13^. In photosynthetic organisms, the positioning of photosynthetic machinery plays a key role in adapting to the environment. For example, in plant leaves, actin-driven re-positioning of chloroplasts between the light-exposed cell surface and light-sheltered lateral edges of cells maximises light access under low light conditions or mitigates photo-damage under increased light intensity respectively^14–16^. Intriguingly, light microenvironments exist within coral tissues that can vary by up to tenfold depending on the tissue orientation with respect to the light and local tissue morphology^17–19^. Cnidarian polyps have been shown to house symbionts in different abundances across anatomical compartments, including the tentacles and oral disk at the oral end of the polyp, and body column and pedal disk at the aboral end^20–24^. Typically, tissues at the oral surface of polyps receive maximal exposure, with light attenuating towards the aboral tissues^25,26^, indicating that a symbiont’s position along the host aboral-oral body axis determines its light access for photosynthesis. These studies together imply that the distribution of symbionts within the host likely plays a key role in the environmental response of symbiosis. How symbiont organisation is regulated within host tissues remains unclear.

Here, we use polyp morphogenesis in the symbiotic sea anemone Aiptasia as a system to study how symbionts become organised in the host concomitant with the emergence of polyp body architecture. We show that an initial endogenous spatial bias of symbionts to the tentacle buds emerges during polyp morphogenesis and that host cells occupied by a symbiont are constrained within the epithelium, consistent with mechanical deformation of host cells by resident symbionts. We observe dynamic expulsion and re-uptake of symbionts from the host gastric cavity during morphogenesis and find that inert spheres grafted into early-stage host tissue lacerates recapitulate the spatial organisation of symbionts along the aboral-oral axis. Finally, we show that symbiont spatial organisation is characteristically remodelled under perturbed light conditions in adult *Pocillopora damicornis* coral polyps. These findings point to a model in which symbionts are patterned at the tissue level by cycles of expulsion into the gastric cavity and re-uptake into host cells at new tissue sites. This opens new questions on the molecular players that determine these dynamics, environmental sensing and responding mechanisms, and the role of symbiont spatial reorganisation in adapting to environmental stress.

## Results

### Symbionts become enriched in the tentacle endoderm during polyp morphogenesis

To investigate how symbionts are spatially organised during polyp morphogenesis, we used a tissue lacerate system in the sea anemone Aiptasia^27^, which forms a symbiosis with the same algal partners as corals but exists as a more experimentally manipulable solitary polyp^28^. Small tissue fragments excised from the base of adult Aiptasia polyps are capable of developing into a new polyp, providing a tractable system for observing symbiont integration into the host (Figure 1A). To test whether an endogenous symbiont organisation emerges during morphogenesis, we cultured tissue lacerates in dark conditions. Following excision, tissue fragments rounded and formed a gastric cavity. At around 5 days, tentacle buds started to form at the oral end concomitant with the formation of an oral opening. The formation of the aboral-oral axis was accompanied by remodelling of actin, with a transition from disordered actin fibres in early, round stages, to circumferentially oriented actin fibres encircling the body column and longitudinally oriented fibres extending along the length of the tentacle endoderm in tentacle budding stages (Figure S1A). The emergence of tentacle buds was accompanied by radial segmentation, similar to that observed in morphogenesis of the non-symbiotic sea anemone Nematostella^29,30^ with actin-rich mesenteries demarcating segment boundaries and segments extending from an aboral base and ending in a single tentacle tip at the oral end (Figure 1B and S1A). These segments provide a modular partitioning of the polyp into individual units with their own aboral-oral axis and symbiont distribution. We imaged segments in tentacle bud-stage lacerates, segmented symbionts and host tissue in 3D, and developed a quantitative framework to map symbiont organisation along the host body axis (Figure S1B, see Methods). We fit a spline through the segment volume from the aboral base to the oral tip and mapped the 3D location of individual symbionts onto the spline (Figure 1C). To investigate how symbionts are distributed within the host tissue, we binned symbiont counts along a normalised aboral-oral axis for each segment, allowing comparison of symbiont organisation across segments. Because the amount of tissue surface available varies along this axis, raw symbiont counts partly reflect how much tissue is present rather than any positional preference. To account for this, we divided the proportion of symbionts in each bin by the proportion of host tissue surface area in that bin, yielding a dimensionless enrichment profile in which a value of 1 indicates symbionts distributed in proportion to the available tissue surface, and values above or below 1 indicating local enrichment or depletion respectively (Figure 1D and S1C, see Methods). This revealed a peak in symbiont enrichment within the oral region of the axis, and a second smaller peak in the aboral region (Figure 1D and S1C). Host cells appear to have a finite capacity to accommodate symbionts^31,32^, suggesting the degree to which a segment is packed with symbionts may shape symbiont distribution along the axis. To explore how symbiont distribution varies with the overall density of symbionts in a segment, we split segments into lower and higher symbiont density groups (total symbiont number divided by total segment surface area, see Methods). This revealed two distinct symbiont enrichment profiles, with less dense segments showing a single peak in symbiont enrichment at the oral end of the axis, whilst more symbiont-dense segments displayed a more uniform symbiont distribution with small peaks in the aboral and oral regions (Figure 1E). To determine whether symbiont organisation was associated with distinct morphological compartments along the aboral-oral axis, we used the circumferential to longitudinal transition in actin to demarcate the transition point from the body column to the tentacle buds and assessed symbiont enrichment across this boundary (Figure 1F, S1A). In less symbiont-dense segments symbionts were enriched in the tentacle bud endoderm versus the body column endoderm, whilst this bias disappeared in host segments with higher symbiont densities (Figure 1G, n = 23 segments across 14 low-density lacerates, 522 symbionts, p = 7.2×10^-7^; n = 13 segments across 9 high-density lacerates, 1592 symbionts, p = 0.27). Together these findings reveal a spatial organisation of symbionts that emerges along the host aboral-oral axis during polyp morphogenesis, with symbionts enriching in the tentacle bud endoderm when overall symbiont density is low. Unexpectedly, this oral enrichment occurs in the absence of an overhead light source. This suggests that symbionts are preferentially patterned to the tentacle endoderm - linking symbiont spatial organisation to tentacle morphogenesis.

### Endosymbiotic host cells exhibit constrained motion within the host epithelium

To explore mechanisms that could spatially organise symbionts along the aboral-oral axis, we live-imaged and tracked symbiont-occupied cell movement during Aiptasia morphogenesis, taking advantage of the autofluorescence of the algal symbionts (Movie S1). Cells were tracked in clusters (tracks in 2D on maximum intensity projections of z-stacks) and the bulk motion of the tissue (see Figure S2) was removed to obtain the relative motion of cells within each cluster. Symbiont-occupied cells displayed constrained motion within the host epithelium over a period of ∼5 hours (Figure 2A and B, MSD α<1. n=4 lacerates, 51 cell tracks). Constrained cell movement can result from packing cells with large organelles, such as an expanded nucleus^33^. To explore a potential mechanical basis for the limited motility of symbiont-occupied cells, we examined their cell morphology. Using actin cortex and membrane labels to visualise cell outlines in fixed samples, we found that symbiont-occupied host cells were significantly expanded and rounded in shape compared to their unoccupied neighbours (Figure 2C and D, n=9, n=8 cells), with the effective volume fraction of symbionts within host cells (∅_*S*_ ∼2) exceeding values known to induce jamming in other epithelial contexts^33^ (Figure 2C, n=7 symbiont/unoccupied host cell area comparisons, mean volume fraction =2.098±0.84, see Methods). Additionally, unoccupied host cells located between symbiont-occupied cells displayed elongated aspect ratios, indicative of cell compression (Figure 2E and F, n=12 cells in aposymbiotic regions, n=8 between symbiont-occupied cells, p = 0.00054). These findings suggest that the internalisation of large symbionts mechanically deforms host cells and limits their motility in the epithelium. To explore whether global tissue flows move symbiont-occupied cells to the oral region of the axis and tentacle buds, we tracked the aboral and oral displacement of symbiont-occupied cells located either aborally or mid-way along the aboral-oral axis of lacerates over a 5-hour period. Symbiont-occupied cells moved in both the aboral and oral directions, with no evidence of net directed movement along the axis (Figure 2G, mean velocity = 0.0035± 0.0074 fractional units/hr in the aboral direction, n = 3 lacerates, 30 tracks). Thus, neither extensive rearrangement of symbiont-occupied host cells or global tissue flows are likely to drive the spatial organisation of symbionts within host tissues.

**Figure 2.**
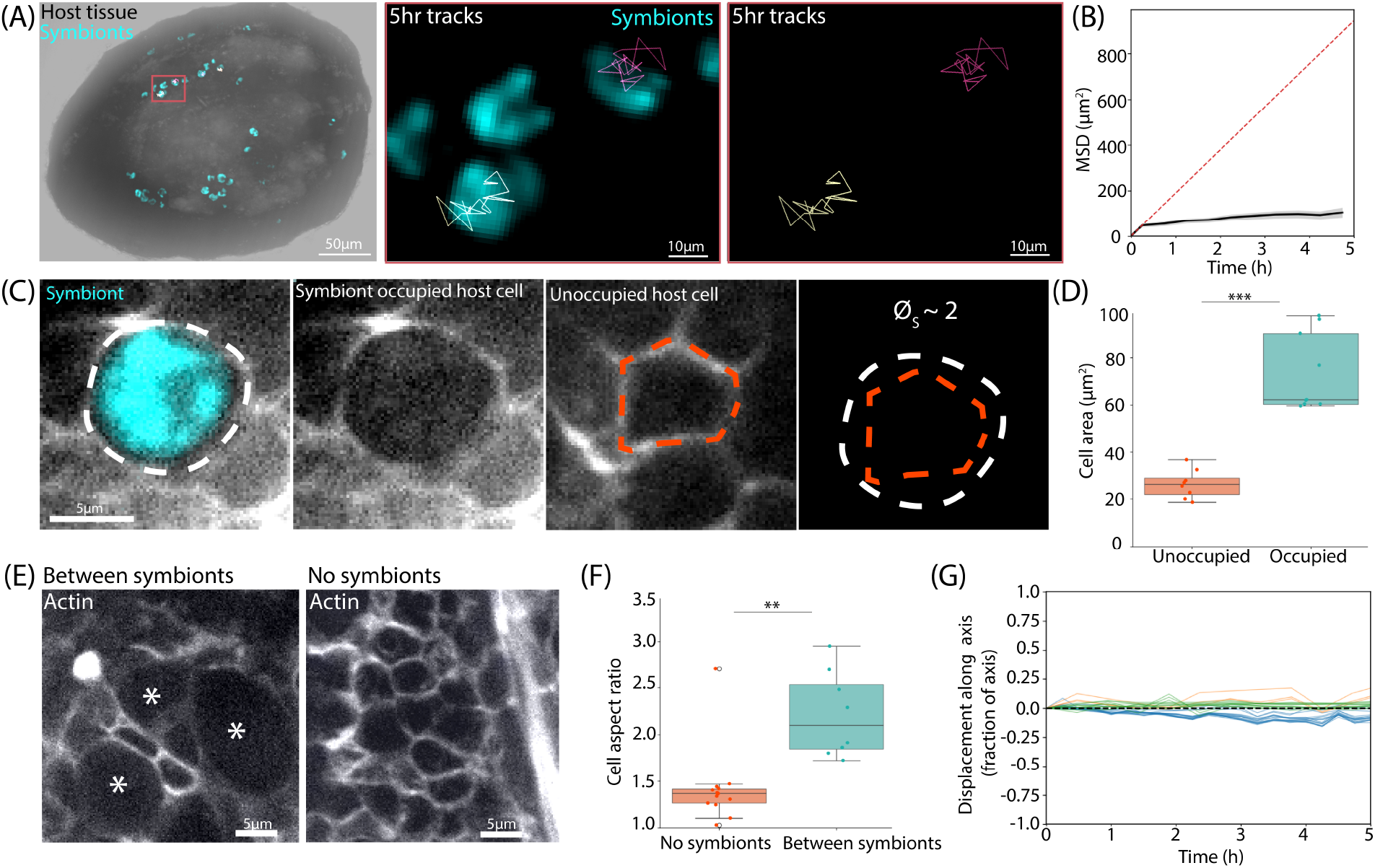
Symbiont-occupied host cells appear mechanically deformed and constrained within the host endoderm. (A) Tracking symbiont-occupied host cell movement during morphogenesis. Symbionts are shown in cyan. Individual cell tracks shown in different colours are visualised in zoomed views of the ROI. (B) Mean squared displacement of symbiont tracks in the host epithelium. The red dashed line indicates diffusive motion, the black line shows the ensemble curve of all tracked symbiont-occupied cells (n=51 tracks). (C) Traced outlines of representative symbiont-occupied (white dashed line) and unoccupied host cells (orange dashed line), using actin (white) to distinguish the cell outline. (D) 2D area (taken in a plane through the approximate mid-point of each cell) of unoccupied and symbiont-occupied host cells (n = 8 unoccupied, 9 occupied cells; p=1.52×10^-6^). (E) Representative confocal image of unoccupied host cells located between symbiont-occupied cells (left, symbiont-occupied cells marked by *) and in a region with no symbionts (right). Actin (white) is used to visualise cell outlines. (F) Aspect ratio of host cells in regions without symbionts and in regions between symbiont-occupied cells (n=12 cells in aposymbiotic regions, n=8 between symbiont-occupied cells; p = 0.00054). (G) Relative displacement of symbiont-occupied cell tracks along the aboral-oral axis (1 corresponds to a full aboral-oral axis length).

### Dynamic cycles of expulsion and re-uptake translocate symbionts within the host

In our tracking analysis, we observed the disappearance of a subset of symbionts between timeframes, suggesting that they may have been expelled from the host cells they occupied. In Aiptasia larvae, dynamic expulsion and re-acquisition of microalgae occurs under conditions where a stable endosymbiosis fails to develop - with microalgae being vomocytosed from the host cell and expelled back into the environment through the larval mouth^34^. Imaging tissue lacerates at a higher framerate (100ms frame interval) confirmed that symbionts exited the host epithelium and entered the fluid-filled gastric cavity where they exhibited relatively rapid motion (Figure S3A, Movie S2). Extended timelapses over a 15-hour period captured repeated symbiont expulsion from the host epithelium and uptake at different tissue locations (Figure 3A, Movies S3-S5; n = 9 expulsion, 12 uptake events across ∼159 trackable symbionts in 4 lacerates), indicating that a fraction of symbionts undergo positional change on a timescale of hours through dynamic expulsion and re-uptake (mean percentage expulsion events per sample = 6.6%, mean percentage uptake events per sample = 10.6% within 15 hours). To explore whether the same symbiont that was expelled was re-acquired at a different location, we attempted to follow the path of individual symbionts. In a small number of cases, we were able to track symbionts in 3D as they exited the host endoderm, moved rapidly within the fluid-filled cavity, and reemerged in a new tissue location (Figure 3B, S3B and S3C, Movie S6). These sequences demonstrate that individual symbionts can relocate along the aboral-oral axis via the gastric cavity (n=4/4 relocations in the oral direction). Whilst the number of fully tracked events is low, these observations suggest that dynamic symbiont expulsion into- and re-uptake from-the gastric cavity may provide a mechanism to spatially organise symbionts along the aboral-oral axis over a timescale of days in which polyp morphogenesis takes place.

**Figure 3.**
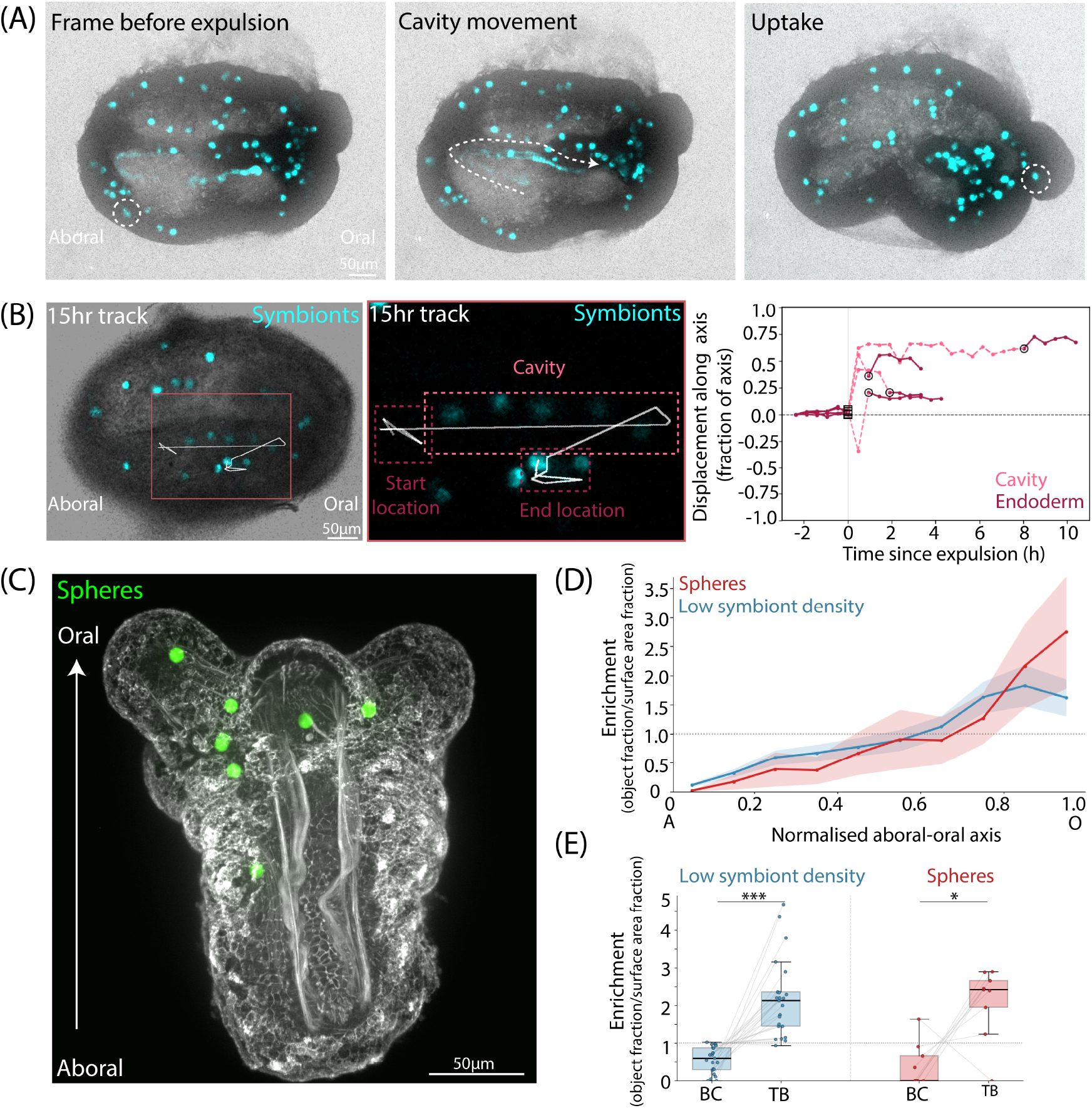
Live imaging of symbiont movement during host morphogenesis and investigation of inert sphere distribution along the host aboral-oral axis. (A) Montage showing a symbiont expulsion event, movement of a symbiont within the host gastric cavity, and an uptake event into a tentacle bud. (B) Overview and ROI images of a continuous track following a symbiont through expulsion, movement within the gastric cavity and re-uptake into the host endoderm. The accompanying plot shows all four trackable cases of symbiont expulsion and re-uptake and the corresponding displacement along the axis. (C) Representative image of polystyrene spheres (green) in a tentacle bud stage aposymbiotic lacerate following morphogenesis. (D) Pooled enrichment profile of spheres (red, n=15 segments from 9 lacerates, 28 spheres) and symbionts from low-density segments from Figure 1 (blue, n=23 segments from 14 lacerates, 522 symbionts) along aboral-oral axis. (E) Symbiont enrichment in the body column (BC) vs tentacle bud (TB) region of segments of low symbiont density segments from Figure 1 (n=23 segments, p=7.2×10^-7^) and sphere segments (n=9 segments from 7 lacerates, 22 spheres, p=0.02).

At the cell level, symbiont expulsion and uptake are associated with different mechanisms of regulation, with host-symbiont interactions underpinning the decision to expel a symbiont whilst the initial uptake of a symbiont by a host cell has been shown to be predominantly indiscriminate, with endoderm cells readily internalising inert spheres^22,34^. To test whether the spatial bias in symbionts to the host tentacle buds was regulated by a host-intrinsic mechanism or host-symbiont signalling, we grafted fluorescent polystyrene spheres with a diameter matching that of symbionts (∼7µm) into aposymbiotic (symbiont-lacking) tissue lacerates. Spheres became internalised into the endoderm during morphogenesis and transiently displayed rapid motion indicative of sphere uptake and subsequent expulsion into the gastric cavity. We assessed the distribution of spheres along the aboral-oral axis in tentacle budding stage lacerates (Figure 3C-D). Intriguingly, the sphere distribution along the aboral-oral axis of tissue segments was consistent with that of symbionts from low symbiont-density segments in our previous experiments (Figure 3D). Whilst sphere number per segment was low, internalised spheres enriched more in the tentacle bud endoderm compared to the body column (Figure 3E, n=9 segments across 7 sphere lacerates, 22 spheres, p=0.02), suggesting that symbiont spatial organisation during host morphogenesis is regulated by a host-intrinsic mechanism at the level of cell uptake.

### Light perturbations induce spatial reorganisation of symbionts in coral polyps

Our data point to expulsion and subsequent re-uptake as a candidate mechanism to spatially organise symbionts during host morphogenesis. As environmental stress also drives symbiont expulsion, we hypothesised that control of spatial organisation can act as a responsive/adaptive mechanism to environmental change. To test this, we leveraged a reef-building coral, *Pocillopora damicornis*, that is composed of stationary polyps amenable to both long-term environmental perturbation and visualisation of symbiont organisation and increased the light intensity incident from directly above from an acclimatised PPFD (Photosynthetic Photon Flux Density) of 25 µmol m^-2^s^-1^ to 50µmol m^-2^s^-1^. We observed a pronounced change in symbiont distribution over the course of 18 days (Figure 4A). The symbiont fluorescent signal intensity within individual segments along the aboral-oral axis showed enrichment in the body column and clearing of the mid region of tentacles (Figure 4B-C). In addition, distinct symbiont clusters appeared along the lateral edges of tentacles at 4 days post light increase, and became more resolved as time progressed (Figure 4C-D, n=5 day 0, n=4 +4 days, n=5 +18 days). These observations show that the tissue-level symbiont organisation in host polyps is responsive to changes in the light environment.

**Figure 4.**
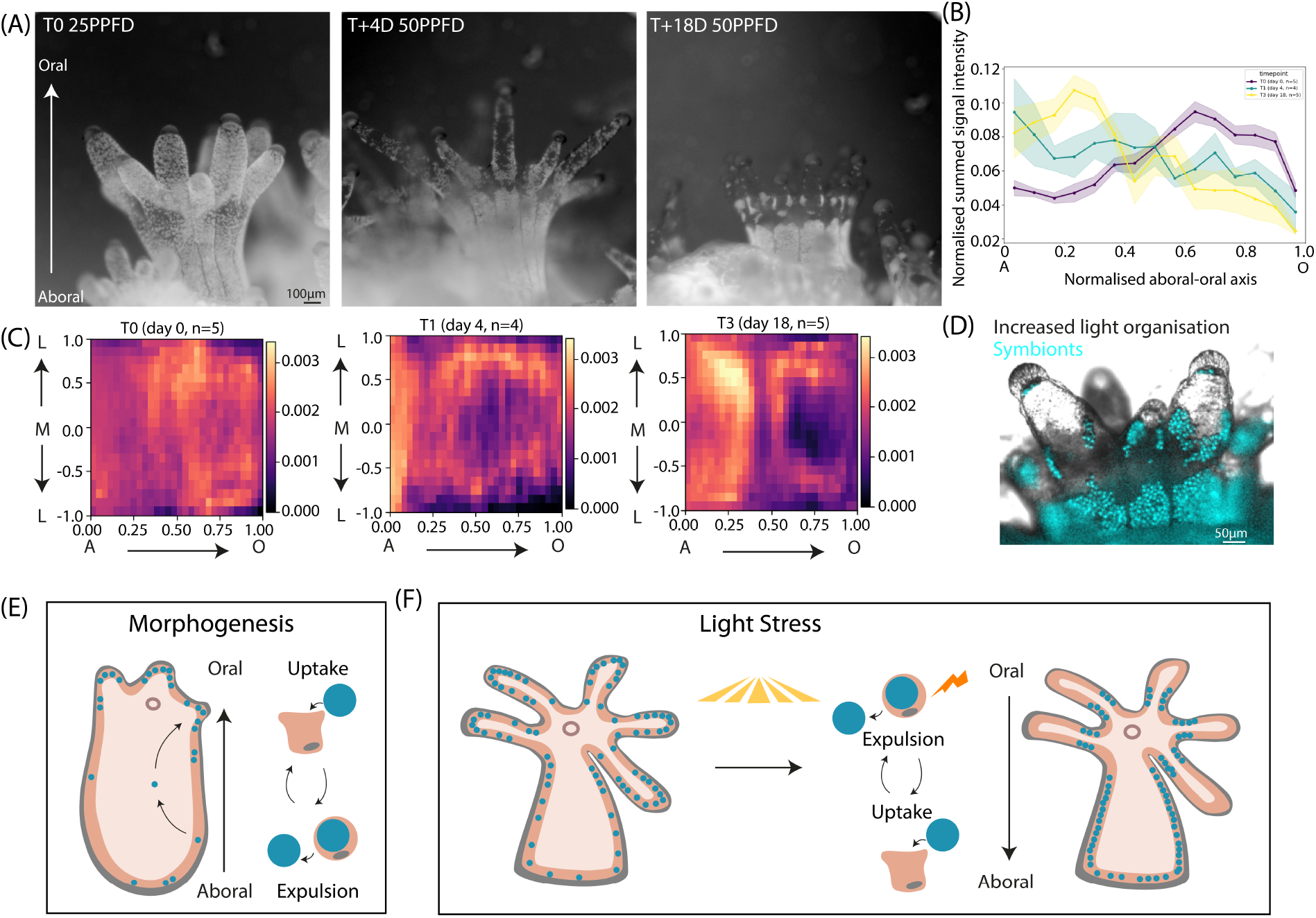
Symbiont organisation in host tissues is altered under light perturbation. (A) Inverted brightfield images showing symbiont organisation in Pocillopora damicornis coral polyps at T0 (acclimatised light, 25PPFD), T +4 days at 50PPFD, T +18 days at 50PPFD. Symbionts appear as white dots. (B) Symbiont signal intensity (bright pixels) along the aboral-oral axis of polyp segments. (C) Averaged heatmap of signal intensity across segments per timepoint. Each segment is unwrapped such that the lateral extents of the segment project to 1 and -1 on the Y axis, and the aboral and oral extents to 0 and 1 on the X axis. (D) Widefield image showing symbiont organisation in a polyp post light-increase. Symbionts are shown in cyan. (E-F) Proposed mechanism of symbiont spatial organisation within the host. (E) A host-intrinsic bias in symbiont uptake, acting within regional cycles of expulsion and re-uptake along the aboral-oral axis, may enrich symbionts in the tentacle buds during morphogenesis. (F) Altered light may shift the regional balance of expulsion and re-uptake, remodelling symbiont organisation within the host.

## Discussion

The spatial organisation of specialised cells plays a fundamental role in development and adaptation across multicellular systems^35–37^. Here, we show that the spatial organisation of algal symbionts within cnidarian host tissues is a regulated property of symbiosis. An aboral-oral organisation of symbionts emerges in Aiptasia polyp morphogenesis, enriching symbionts to the tentacle bud endoderm. Under altered light conditions, symbiont organisation is remodelled in adult *Pocillopora damicornis* polyps, shifting symbiont distribution both along the aboral-oral axis of host segments and laterally in the tentacles. The establishment of symbiont organisation in morphogenesis occurs without the rearrangement of symbiont-occupied host cells, which appear mechanically constrained within the host epithelium, linking the symbiotic state of host tissues to tissue mechanics and contrasting with the extensive cell rearrangements documented in a non-symbiotic cnidarian^6^. Instead, dynamic cycles of symbiont expulsion into, and re-uptake from the fluid-filled host gastric cavity enable the long-range translocation of symbionts within the host. Spatially organising symbionts through expulsion and re-uptake via a fluid cavity may provide a mechanism to accommodate large symbionts in an epithelial tissue context, relaxing/unjamming the host tissue under a potential mechanical constraint. Our sphere grafting experiments support a host-intrinsic mechanism that establishes tentacle-biased symbiont enrichment during morphogenesis and point to regional differences in symbiont uptake as a point of regulation (Figure 4E). Such differences may be pre-patterned by host cell capacity to phagocytose large symbionts, for example through regional differences in host cell cortex tension posing a mechanical gate on phagocytosis^38^. Whether symbiont reorganisation under perturbed light conditions is driven by a host-intrinsic mechanism or by signals from the symbiont remains to be determined. We hypothesise that stress-altered dynamics of symbiont expulsion and re-uptake remodel symbiont organisation under environmental change (Figure 4F). A symbiont’s position within the host is likely to set the light conditions it experiences, as scalar irradiance varies by several-fold along the aboral-oral axis of coral tissue^17,25,26^ across a steeply structured internal light field^19^. Cells in tissue regions that experience higher light exposure may decrease symbiont uptake or increase symbiont expulsion, shifting symbiont localisation toward more sheltered tissue locations. A symbiont-derived signal could come from photoinhibition of the symbiont photosystem under excess light, driving higher levels of ROS production and a decline in photosynthate in the most illuminated tissue regions^8,9^; such a spatially graded signal could locally raise symbiont expulsion through immune-triggered vomocytosis^34^. Alternatively, host cells may directly sense light intensity, for example via a light-responsive opsin known to be expressed in symbiont-hosting cells^39^.

The endogenous enrichment of symbionts to the tentacle endoderm during morphogenesis and their light-driven clearing from it act on the same, most light-exposed tissue. We propose that this tentacle niche is favourable for light capture when light is limiting and metabolic demands are high, as for a juvenile polyp starting life on the reef floor^27,40,41^ but becomes a liability under excess light, when symbionts are repositioned away from it. The same positional axis is thus read in opposite directions under the two regimes, consistent with the reduced symbiont density observed at the seasonally light exposed tentacle tips of adult corals^21^. The clustering of symbionts to lateral edges of the tentacles is reminiscent of the photoprotective repositioning of chloroplasts to the lateral edges of plant cells^14^. In coral-algae symbiosis, the repositioning of symbionts at the tissue level could place them in protective light microniches, potentially contributing to the lower bleaching susceptibility of thicker-tissue corals which may have a greater capacity to shelter symbionts from excess light^42^. The lateral clustering of symbionts in the tentacles further points to finely resolved light microniches, potentially shaped by spatially patterned host pigments^21,43^. Future studies are needed to uncover the role of symbiont reorganisation in preserving photosynthesis efficiency and buffering environmental stress. By capturing symbiont spatial organisation within cnidarian tissues, our work sets up a quantitative framework for investigating how symbiosis develops and responds to a dynamic environment. Future work using spatial organisation as an emergent, quantitative readout of cellular decisions will help elucidate cellular mechanisms that coordinate the development and response of symbiosis to environmental change.

## Limitations of the Study

The number of inert spheres that successfully internalised into cells of aposymbiotic tissue lacerates in our experiments was low (consistent with previous studies^4^), limiting our ability to compare their distribution with the symbiont distribution profiles at similar levels of particle density. The number of fully tracked expulsion and uptake events is low because unambiguous tracking was only possible when an expelled symbiont did not overlap with other expelled symbionts in the host gastric cavity, and such events were rare. Tracking multiple rapidly moving individual symbionts in the host gastric cavity would require higher frequency imaging, which impairs host viability. The contractile state of host tissues may bias the apparent distribution of symbionts within the host. Whilst tissue lacerate samples were relaxed with MgCl_2_ prior to fixation to limit the effect of differential tissue contractility impacting our distribution analysis, coral polyps were not relaxed in long term timelapses (to support long term sample health and limit confounding effects). Live imaging was performed in filtered ASW in the absence of MgCl_2_ or additional symbionts.

## Acknowledgements

We thank members of the Xiong, Branson, Waller, Gallop, Kawaguchi and Guse labs for technical assistance and constructive feedback. We thank Ben Jenkins, Annika Guse, Buzz Baum and Andrea Dimitracopoulos for feedback on the manuscript. Experimental Aiptasia polyps were originally obtained as a kind gift from Ben Jenkins and the Waller laboratory. We thank Nicola Lawrence and Purnima Kumar from the Gurdon Institute Imaging Facility for microscopy support, Charles Bradshaw from the Gurdon Institute High-Performance Computing Facility for computing support, and the Media Team at the Gurdon Institute for support with all things media. We thank Euan Smithers and the Sainsbury Laboratory for access to a light meter and Jenny Richens and the St Johnston lab for support with custom imaging. **Funding**: this work was supported by a Wellcome Trust/Royal Society Sir Henry Dale Fellowship (215439/Z/19/Z) and UKRI-EPSRC Frontier Research Grant (EP/X023761/1, originally selected as an ERC Starting Grant) to F.X. and a Wellcome/UKRI Physics of Life Grant and Schmidt Sciences Cambridge Centre for Data Driven Discovery and Accelerate Programme for Scientific Discovery Grant to S.B.P.M.

## Author Contributions

S.B.P.M. conceived the project, carried out all experiments and analysed the data. A.J. carried out experiments, data analysis, and maintained experimental cultures. E.S.A. created the custom imaging chamber and carried out spinning disk modification and imaging. X.L. and M.G. carried out exploratory experiments and optimised protocols. D.S.S. and A.B. cultured coral colonies and created coral frags for experiments. F.X. acquired funding and contributed to experimental design and conceptualisation. S.B.P.M. supervised the work, acquired funding, and wrote the manuscript with input from all authors.

## Methods

### Aiptasia Culture

*Exaiptasia diaphana* (commonly termed Aiptasia) of the clonal line CC7, hosting the native dinoflagellate symbiont *Breviolum minutum* (strain SSB01), were maintained in laboratory incubators. Anemones were housed in an incubator in groups of 5-10 at 26 °C under a 12 h light (25PPFD) - dark cycle in 250-500 mL plastic containers containing 33 ppt artificial seawater (ASW). Light conditions created by LED arrays were calibrated using a Licor180 light meter. Polyps were fed *Artemia* nauplii twice weekly during the light phase, and ASW was replaced the following day. Asexual reproduction enabled the continuous production of new individuals for experimental use.

### Coral Culture and Generation of Frags

*Pocillopora damicornis* colonies were originally obtained from Tropical Marine Centre, Chorleywood, UK, and were cultured for ∼ 1 year in 900L recirculating reef aquaria tanks at 26 °C in ASW (Reef Salt TMC ©). Coral frags were created using a circular saw to cut off the tip of a coral branch. Tips were glued down onto glass slides using standard cyanoacrylate super glue (Gorilla Super Glue Gel) and were returned to the main tank for a few days of recovery before moving to the incubator for experimentation. Once in the lab incubator coral frags were cultured under the same conditions as Aiptasia polyps (described above), with the exception that ASW was prefiltered for coral frag cultures.

### Generation of Pedal Lacerates

To maximise comparability across experiments and tractability for analysis, adult Aiptasia polyps were either partially or fully bleached to obtain reduced symbiont density and aposymbiotic polyps that were used for symbiont and sphere organisation experiments. This was achieved through repeated cold shock, where polyps were transferred to 6-well plates and ASW was replaced with pre-chilled (5 °C) filtered ASW (FASW). Plates were incubated at 5 °C for 4-6 h, after which cold FASW was replaced with 26 °C FASW and polyps were returned to the incubator. Polyps were assessed for symbiont presence under a dissection microscope, separated into reduced and aposymbiotic groups, and cultured in dark conditions before use in experiments by placing culture plates on the top shelf of an incubator above the level of the LED arrays to minimise light exposure. Reduced symbiont density polyps were used to generate lacerates that were amenable for segmentation and tracking experiments, whilst aposymbiotic polyps were used for sphere grafting experiments.

Pedal lacerates were generated by placing medium-to-large polyps in a small dish and allowing them to firmly adhere to the substrate. Thin sections (∼1 mm diameter) of the pedal disk were excised using a sterile scalpel and further subdivided with perpendicular cuts. Lacerates were washed by repeated transfer into fresh FASW. For microsphere grafting experiments, freshly prepared lacerates were immediately pressed into a droplet of green fluorescent 7 µm FluoSpheres® polystyrene microspheres using fine tweezers or an eyelash tool. Lacerates were transferred to individual wells of multi-well plates containing FASW and maintained at 26 °C. Lacerates typically initiated budding between 5-8 days post-laceration.

### Fixation and Immunostaining

Tissue lacerates were fixed at early rounded and tentacle budding stages of morphogenesis. Lacerates were imaged daily, and individuals were selected based on overall morphology, tissue integrity, and absence of overt tissue damage. Samples were fixed in 4% paraformaldehyde (PFA) in 1× phosphate-buffered saline (PBS) for 30 min at room temperature in a fume hood, followed by three 30-min washes in PBS containing 0.5% Triton X-100 (PBST) at room temperature with gentle rocking and an additional overnight wash at 4 °C. Samples were then washed three times in PBST and incubated with phalloidin-Alexa Fluor 647 and Hoechst nuclear stain (both 1:200) for one day and overnight at 4 °C. Final washes included three 30-min washes in PBST followed by two 30-min washes in PBS. Samples were mounted in domed droplets of VectaShield on glass-bottom dishes and stored at 4 °C until imaging.

### Custom Imaging Chamber and Confocal Spinning Disk Live Imaging

Lacerates were imaged in a custom holder that incorporated interchangeable standard 35 mm diameter cover glass bottom dishes (Ibidi, 81218-200) into a sealed flow chamber. Samples were grown in an environmentally controlled incubator as described above. As the lacerates matured, just prior to tentacle formation, the 35 mm dishes they were adhered to were transferred to the custom chamber without the need to remove and transfer the lacerates. That is, they were maintained in a 35 mm cover glass bottom dish without disruption during growth and imaging. The custom chamber allowed a 35 mm dish to be placed on the microscope sample XYZ stage. The dish lid was replaced with an o-ring sealed lid that included two luer port connections for fluid flow as shown in Supplemental Figure S3D. The mounting arrangement also securely held the dish for XYZ positioning. Once a 35 mm dish was mounted on the microscope, tubing was connected to the luer ports. One tube was passed through a peristaltic pump (Ismatec, REGLO ICC ISM4412). The ends of both tubes were placed in a conical flask containing approximately 3 Liters of FASW (filtered artificial seawater). The seawater flask was maintained at 26°C via a heating pad (Vivosun, 330101) and the peristaltic pump was set for a flow rate of approximately 100 µl/min. When the pump was switched on, FASW from the flask was gently circulated through the chamber and back into the flask as shown in Supplemental Figure S3D.

Imaging was performed on a custom spinning disk microscope. In brief, 642 nm excitation laser light (MPB Communications, 2RU-VFL-P-2000-642-B1R) was coupled into a 600 µm core multimode fiber connected to a Crest Optics X-Lite V2 confocal spinning disk with 70 µm pinholes. The X-Lite V2 unit was positioned with the spinning disk at the IX83’s primary image plane. Excitation light was reflected into the common light path with a dichroic mirror (Chroma, 59007bs). Fluorescence emission was collected by a 10X, 0.4NA objective lens (Olympus, UPLXAPO10X), passed back through the spinning disk, was separated from the excitation laser light with the dichroic mirror mentioned above, and further separated into a far-red channel with a second dichroic mirror (Chroma, ZT647rdc) and emission filter (Chroma, ET700/75). Fluorescence was captured with an sCMOS camera (Photometrics, Prime BSI) with a 650 nm effective pixel size and a 900 x 900 pixel image format resulting in a 5.85 x 5.85 mm^2^ field of view. A second, bright field, channel was simultaneously collected on a second camera. For this, the sample was Köhler illuminated via the IX83’s transmitted light lamp with the addition of a narrow band filter (Semrock, FF01-600/52) to avoid interference with the far-red channel. Transmitted lamp light followed the same optical path but was separated from fluorescence emission with the previously mentioned dichroic mirror and directed onto a separate sCMOS camera (Photometric, Prime BSI). Microscope control and data collection was carried out with a freely available software in the LabView environment (github.com/Gurdon-Super-Res-Lab/Microscope-Control).

Prior to imaging, candidate samples were visually identified, and each position was marked and saved. After all samples of interest had been identified and marked, the microscope collected a 400 µm z-stack with a 1 µm step size and 100 ms exposure time per step at each marked location. After the collection of z-stacks at each marked position, the microscope paused for 25 minutes before collecting another set of z-stacks at each marked position. Z-stacks were repeated every 25 minutes for 24 to 72 hours.

### Confocal Imaging of Fixed Samples

Fixed pedal lacerates were imaged using a Nikon SoRa spinning-disk confocal microscope to capture host cell morphology, three-dimensional symbiont localisation, and cellular densities across Z-stacks. Brightfield imaging and laser excitation were used as appropriate, including 638 nm excitation to visualise algal symbiont autofluorescence. Imaging was performed using 10× air, 20× water immersion, 40× water immersion, and 60× water immersion objectives. Both this Nikon SoRa spinning-disk confocal and the custom spinning disk imaging set-up described above were used to acquire short-term live imaging for MSD analysis.

### A quantitative framework for mapping symbiont organisation along the host body axis

Confocal stacks were pre-processed slice-by-slice with background-subtraction (Gaussian blur, σ = 20 px), percentile-based intensity normalisation, and light smoothing. Symbiont segmentation was performed in 3D with Cellpose v4.0.7 using the pretrained Cellpose-SAM model (cpsam^44^) on GPU. Each tissue segment underwent manual quality control; only segments in which individual symbionts were clearly resolved were retained, and symbiont masks were filtered by equivalent-sphere diameter (≥ 5 µm) and 3-D sphericity (≥ 0.6) to remove spurious labels.

#### Tissue segment masks

Anatomically distinct tissue segments demarcated by actin-rich boundaries were outlined interactively in napari as 2-D polygons on transverse cross-sections; a 3-D mask was built by linearly interpolating the polygons’ signed-distance transforms along Z, and adjacent segments were separated by watershed (σ = 2.0, compactness = 0) using Sobel edges of the Gaussian-filtered image as the landscape. Each voxel was assigned to at most one segment.

Where needed, volumes were rotated to align the aboral-oral axis to a standard view (orthonormal basis from the napari camera) and resampled to isotropic voxels at the minimum input spacing (typically 0.18-0.3 µm; scipy.ndimage.affine_transform, linear for intensity images, nearest-neighbour for labels).

#### Aboral-oral axis and spline fitting

For each segment the aboral-oral axis was seeded by two anchor points (aboral base, oral tip) and a centroidal spline was fitted iteratively (2 passes) through the centre of mass of segment cross-sections. Up to 500 segment voxels were projected onto the current axis and binned into 10 equal cross-sectional slices, bin centroids were smoothed (1-D Gaussian, σ = 3 bins), and a parametric cubic spline (scipy.interpolate.splprep, s = 0) was fitted through them anchored at both endpoints. Each reconstructed segment was reviewed manually in napari, where a reviewer recorded a pass/fail verdict for the symbiont mask (whether individual symbionts were clearly resolved), the tissue-segment mask and the aboral-oral spline fit. A segment was retained for further analysis only if all checks passed.

#### Symbiont and sphere projection onto the aboral-oral axis and enrichment analysis

Each symbiont centroid (for symbiont distribution analysis) or sphere centroid (for inert sphere analysis) was projected onto its segment’s spline at the nearest arc-length point and given a normalised aboral-oral position on [0, 1] (0 = aboral, 1 = oral), allowing symbiont distributions to be compared across segments of different aboral-oral axis lengths. Examples of symbiont counts binned along the raw aboral-oral axis are shown in Figure S1D.

To account for the amount of host tissue available at each position, and to allow comparison across segments and groups with different overall symbiont loads, symbiont/sphere distribution was expressed as an *enrichment*: the proportion (number divided by total amount) of objects in a bin divided by the proportion of host tissue surface area in that bin (a value of 1 indicates objects track available tissue).

For individual segment profiles (Figure S1C) the enrichment of bin *i* in segment *k* was calculated as

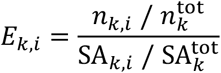

where *n*_*k,i*_ and SA_*k,i*_ are the object count and tissue surface area in bin *i* of segment *k*, and 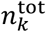 and 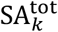 are that segment’s total object count and total surface area.

For pooled profiles (Figure 1D and E, Figure 3D) the enrichment of bin *i* was calculated as

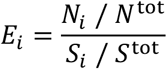

where *N*_*i*_ and *S*_*i*_ are the object count and tissue surface area in bin *i* summed across all segments, and *N*^tot^ and *S*^tot^ are the totals across all segments. For body column and tentacle bud compartment comparisons (Figure 1G, Figure 3E), enrichment is each compartment’s proportion of a segment’s symbionts divided by the compartment’s proportion of that segment’s tissue surface area.

Host tissue surface area was measured as the area of a marching-cubes mesh of the segment, resampled to isotropic 1 µm voxels to improve standardisation of the analysis across samples that were imaged with different objectives and voxel sizes due to sample size variation. Enrichment was profiled over 10 bins along the normalised axis and smoothed (Gaussian, σ = 0.8 bins). For each bin position, the mean and standard error of the mean (SEM) were calculated across segments. Analyses were performed using NumPy, pandas, scikit-image, and matplotlib. We used the circumferential-to-longitudinal transition in actin fibre alignment to mark the body column-to-tentacle transition (Figure S1A) when comparing body column and tentacle bud tissue regions. Only segments with ≥2 objects were included in boxplot comparisons.

#### Segment symbiont density grouping

To assess whether the symbiont distribution depended on overall symbiont load within a segment, each segment’s overall symbiont density (total symbiont number divided by total segment surface area) was computed, and segments at or above the mean density were assigned to the higher density group and the remainder to the low density group.

#### Statistics

Symbiont enrichment in the body-column versus tentacle-bud endoderm was compared using a paired Wilcoxon signed-rank test (scipy.stats.wilcoxon), with each segment contributing paired body-column and tentacle-bud values. Between 1-3 segments contributed per sample in this analysis. Individual segments, each with its own aboral-oral axis, were used as the unit of replication.

### Symbiont distribution analysis in coral polyps under light perturbation

Brightfield images of *Pocillopora damicornis* polyps were acquired using a Ximea camera fitted to a bench-top dissection scope. For each polyp, a segment was outlined manually as a polygon in napari, and three ordered landmarks - the aboral base, the body-column-tentacle transition and the oral tip - were placed to define the aboral-oral axis. A parametric cubic spline was fitted through the three landmarks (scipy.interpolate.splprep, s = 0) and sampled at 200 points; every pixel was projected to its nearest spline point by arc length, giving a normalised aboral-oral coordinate (0 = aboral, 1 = oral).

Symbiont distribution was approximated from a background-subtracted image-intensity signal. Background was removed per segment by subtracting a planar surface fitted through operator-selected tissue-free points, with near-saturated pixels excluded. For the aboral-oral profiles (Figure 4B), background-subtracted intensity was binned into 15 equal bins along the normalised axis and normalised so that the total signal summed to one per polyp; the mean ± SEM across polyps is shown per timepoint. For the averaged heatmaps (Figure 4C), each segment was unwrapped onto a 30 (aboral-oral) × 20 (cross-axis) grid: aboral-oral position was given by arc length along the spline (0 to 1), and the signed perpendicular offset of each pixel from the axis was normalised, within each aboral-oral bin, to that bin’s local half-width, so that every segment fills the range [−1, 1] on the cross-axis irrespective of its shape. Per-polyp grids (pooling multiple focus views of the same polyp) were normalised to sum to one and averaged across polyps per timepoint.

### Cell and Symbiont Shape Analysis

We used the polygon tool in ImageJ to draw around the outline of cells and symbionts in the same Z plane, approximately at their midpoint. Symbiont volume fraction was obtained by dividing symbiont area by the area of an unoccupied host cell within the same sample, as host cell outlines in symbiont-occupied cells were difficult to distinguish due to extensive cell deformation. T-tests (scipy.stats.ttest_ind) were used to statistically compare cell shapes.

### Live cell tracking and motion analysis

Symbiont-occupied host cells were tracked manually in napari by marking each cell’s position in every frame. Each track was assigned by hand to a cluster corresponding to a tissue region. Tracks were trimmed to their single longest run of consecutive frames (maximum gap = 1 frame), and tracks spanning fewer than two frames were discarded. Movies were maximum-intensity projections and motion was analysed in two dimensions for the within-tissue tracking analysis. To remove bulk tissue motion, a per-frame similarity transformation (rotation, scaling and translation) was fitted to the tracked cells and used to register every frame onto the reference frame (the frame containing the most tracks), with registration performed per cell cluster where possible or otherwise on all tracks. Cell motion was analysed as residual displacement in this tissue reference frame. Time-averaged mean squared displacement (MSD) was computed per track, and the anomalous exponent α was the slope of a linear fit to log(MSD) versus log(lag) over a fixed physical-time window. To quantify displacement along the aboral-oral axis, the oral and aboral extremes of each segment were marked and interpolated between frames. Each cell’s position was expressed as a fraction of the current axis length (0 = aboral, 1 = oral), normalising out tissue contraction. For expulsion and re-uptake tracking analysis tracks were made in 3D using the raw Z-stacks to follow symbiont movement through different Z planes as they exited and re-entered the tissue.

#### Software

All analyses were performed in Python 3.9 using Cellpose^45^ for segmentation, napari^46^ for interactive 3-D annotation, ImageJ for visualisation^47^, scipy and scikit-image for spatial analysis and meshing, and scikit-learn for clustering. Claude Code (Anthropic) was used for coding assistance and debugging, predominantly with the Opus 4.8 model. Coding outputs were checked for accuracy using visual inspection throughout the analysis pipeline and cross-checking with manual analysis.

## Notes

### Competing Interest Statement

The authors have declared no competing interest.

