## Supplementary Information for "Symbiont spatial organisation is dynamically regulated within cnidarian host tissues"

### Supplementary Figures

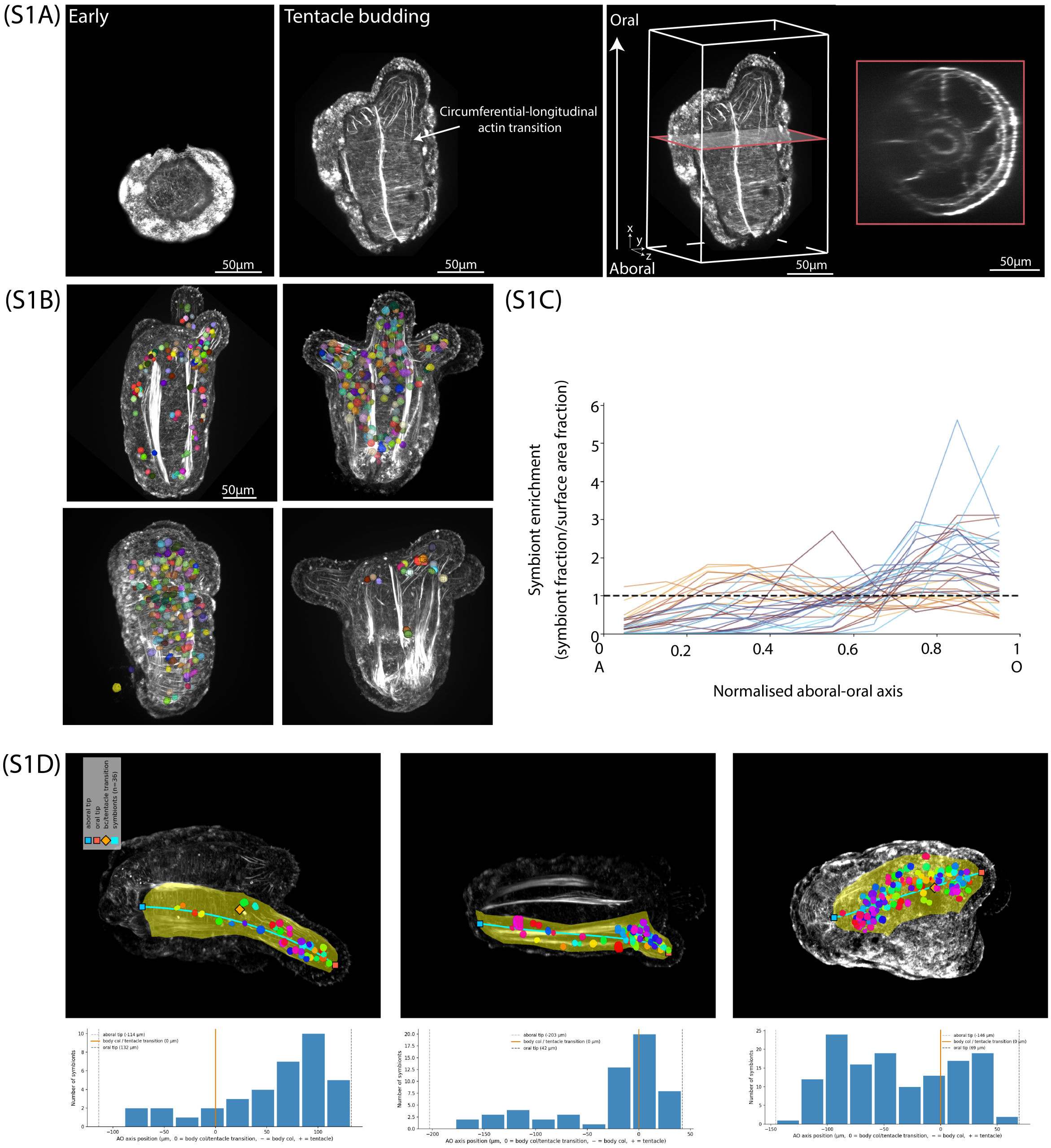


Figure S1. (A) Representative confocal images of actin (white) organisation in early and tentacle budding stage tissue lacerates, and an example view of the host internal structure in cross-section. (B) Example 3D projections of tissue lacerates with symbionts segmented in 3D. (C) Symbiont enrichment profiles for each segment (binned symbiont fraction/surface area fraction) along the normalised aboral-oral axis (n=36 segments from 18 lacerates, 2114 symbionts). Different colours correspond to different segments. (D) Example images of symbiont mapping onto segment splines and corresponding binned symbiont counts along the aboral-oral axis for each segment.

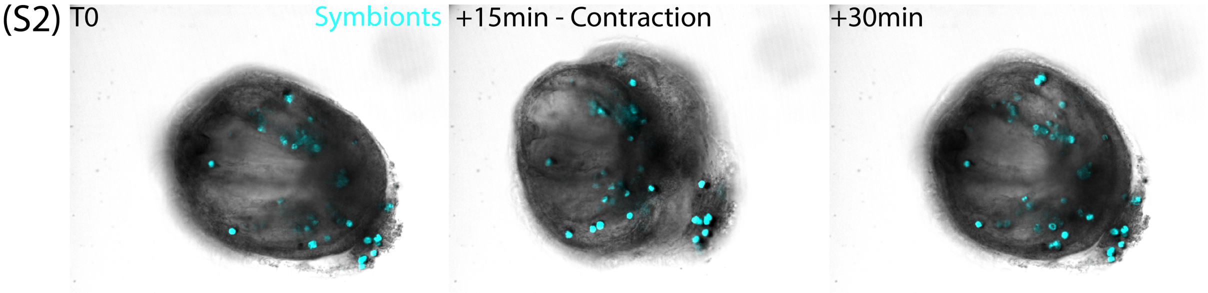


Figure S2. A representative montage showing the pulsatile contractions exhibited by tissue lacerates during morphogenesis. Symbionts are shown in cyan.

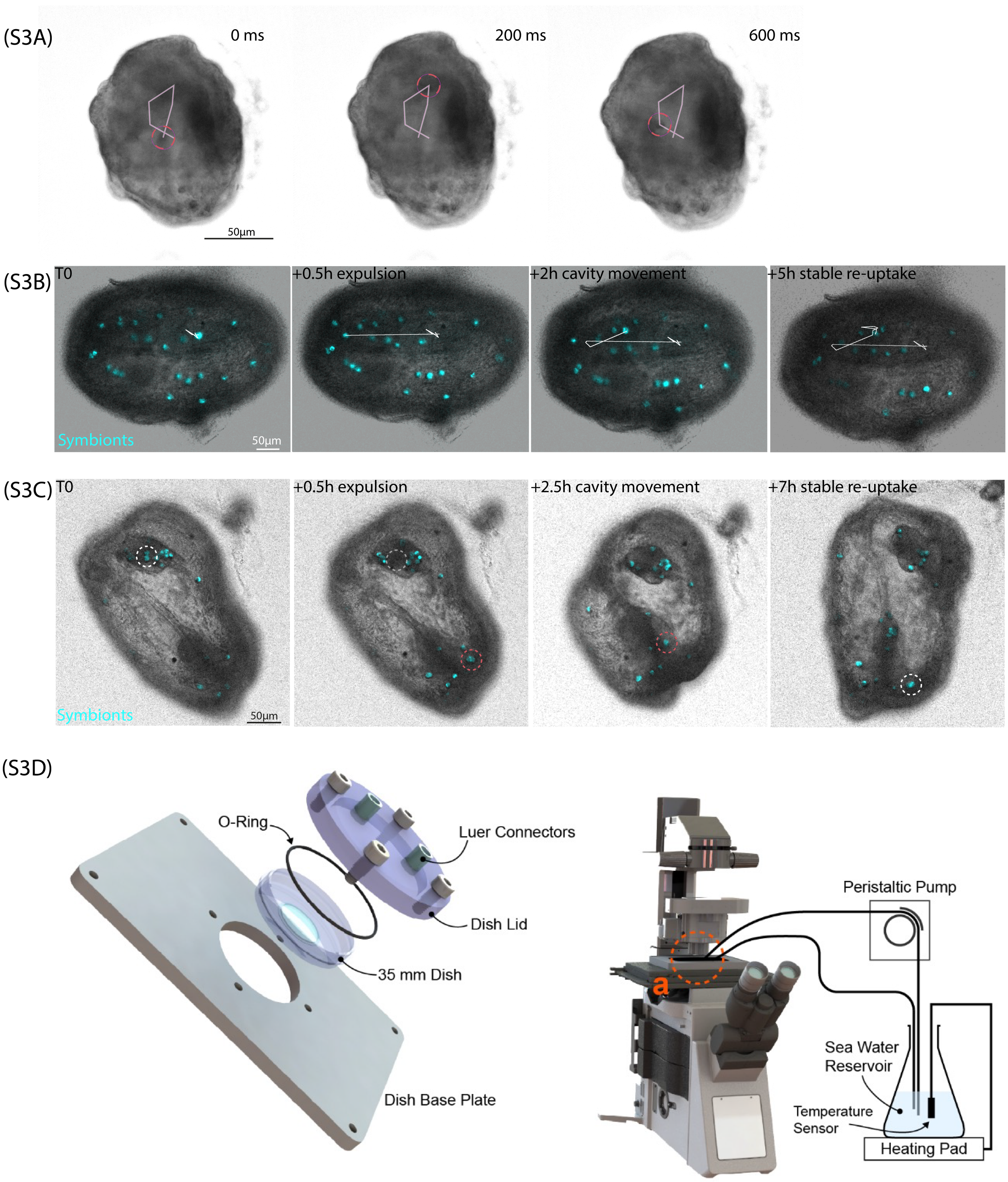


Figure S3. (A) Rapid movement of symbionts within the fluid-filled gastric cavity. The full track is visualised and the position of the symbiont in each frame is annotated by the circle. (B) Montage showing the individual frames corresponding to continuously tracked symbiont expulsion-cavity movement-reuptake. (C) Montage showing an additional example of symbiont tracking throughout expulsion, cavity movement, and re-uptake. (D) Custom imaging chamber and fluid flow. A custom chamber (left) was fabricated from plexiglass and used for sample imaging. The Dish Base Plate was connected to the microscope sample XYZ stage. A standard cover glass bottom 35 mm dish was set in the central aperture of the base plate and sealed with an o-ring and custom Dish Lid. Once sealed, the dish was fully filled with FASW and the Luer Connectors on the lid were fitted with tubing for fluid flow. The custom imaging chamber was installed on a microscope base sample XYZ stage and connected to a peristaltic pump and sea water reservoir for continuous fluid flow and temperature control while imaging (right).

### Supplementary Movies

Movie S1 – Spinning disk confocal imaging of a tissue lacerate undergoing morphogenesis with symbionts integrated. Symbionts are shown in cyan.

Movie S2 – High framerate spinning disk confocal imaging of a tissue lacerate undergoing morphogenesis with the motion of a rapidly moving symbiont in the gastric cavity tracked within the annotated circle. The symbiont can be distinguished by its contrast (appears dark) relative to the host tissue in the brightfield channel.

Movie S3 – Spinning disk confocal imaging of a symbiont expulsion event in a tissue lacerate undergoing morphogenesis. Symbionts are shown in cyan. The white arrow tracks the symbiont position in the host tissue. The magenta arrow tracks the symbiont exit into and movement within the gastric cavity.

Movie S4 – Spinning disk confocal imaging of a symbiont uptake event in a tissue lacerate undergoing morphogenesis. Symbionts are shown in cyan. The magenta arrow tracks the symbiont movement within the gastric cavity. The white arrow tracks the symbiont uptake into the host tissue.

Movie S5 – Long-term spinning disk confocal imaging of symbiont motion and tissue lacerate morphogenesis for the sample shown in Movies S3 and S4. Symbionts are shown in cyan.

Movie S6 – Spinning disk confocal imaging of a tissue lacerate undergoing morphogenesis with a continuously tracked symbiont expulsion and re-uptake event. Symbionts are shown in cyan.
